# tstrait: a quantitative trait simulator for ancestral recombination graphs

**DOI:** 10.1101/2024.03.13.584790

**Authors:** Daiki Tagami, Gertjan Bisschop, Jerome Kelleher

## Abstract

**Summary:** Ancestral recombination graphs (ARGs) encode the ensemble of correlated genealogical trees arising from recombination in a compact and efficient structure, and are of fundamental importance in population and statistical genetics. Recent breakthroughs have made it possible to simulate and infer ARGs at biobank scale, and there is now intense interest in using ARG-based methods across a broad range of applications, particularly in genome-wide association studies (GWAS). Sophisticated methods exist to simulate ARGs using population genetics models, but there is currently no software to simulate quantitative traits directly from these ARGs. To apply existing quantitative trait simulators users must export genotype data, losing important information about ancestral processes and producing prohibitively large files when applied to the biobank-scale datasets currently of interest in GWAS. We present `tstrait`, an open-source Python library to simulate quantitative traits on ARGs, and show how this user-friendly software can quickly simulate phenotypes for biobank-scale datasets on a laptop computer.

**Availability and Implementation:** `tstrait` is available for download on the Python Package Index. Full documentation with examples and workflow templates is available on https://tskit.dev/tstrait/docs/, and the development version is maintained on GitHub (https://github.com/tskit-dev/tstrait).

## Introduction

Genome-wide association studies (GWAS) identify genetic variants that are statistically associated with a specific trait (Uffelmann et al., 2021). Many loci that are associated with various human diseases and traits have been identified (e.g. Yengo et al., 2022; Mathieson et al., 2023), and GWAS results are actively being incorporated into clinical practice (Visscher et al., 2017). The great success of GWAS has prompted the collection of many biobank datasets consisting of hundreds of thousands of participants (Tanjo et al., 2021), but this scale presents significant challenges to current GWAS methodology (Uffelmann et al., 2021).

Simulation is a critical component of GWAS method development, and generally consists of two steps: first simulating genetic variation (genotypes) and then simulating quantitative traits (phenotypes) based on the synthetic genotypes. The combined genotypes and phenotypes represent ground-truth data which GWAS methods can be evaluated against. Genetic variation is usually simulated either by model-based population genetic methods such as `msprime` (Baumdicker et al., 2022) and `SLiM` (Haller and Messer, 2023), or by statistical resampling from existing datasets using methods like `HAPGEN2` (Su et al., 2011) and `HAPNEST` (Wharrie et al., 2023). Both approaches have advantages and disadvantages and excel in different situations. Roughly speaking, model-based simulation methods provide better control of population processes such as demography, whereas resampling methods are better at capturing difficult to model nuances of real data. Model-based population genetic simulations have made great strides in recent years, with major advances in both scalability (Kelleher et al., 2016, 2018; Haller et al., 2018) and realism (Adrion et al., 2020; Anderson-Trocmé et al., 2023), and have been successfully used to simulate large-scale GWAS cohorts (e.g. Martin et al., 2017; Zaidi and Mathieson, 2020).

An important property of these population genetic simulation methods is that they output ancestral recombination graphs (ARGs) rather than sample genotypes. ARGs encode the interwoven paths of genetic inheritance caused by recombination (Hudson, 1983; Griffiths and Marjoram, 1997; Wong et al., 2023), and contain rich detail about ancestral processes. Recent breakthroughs in inferrence methods have made it possible to estimate ARGs at biobank scale (Kelleher et al., 2019; Zhang et al., 2023), and there is now intense interest in their practical application (Lewanski et al., 2024; Brandt et al., 2024). Statistical genetics has been a particular focus, and ARG-based methods have been shown to detect more ultra rare variants than conventional association testing methods (Zhang et al., 2023); to have better power to detect causal loci in quantitative-trait locus mapping (Link et al., 2023); and to provide a sparse and efficient model of linkage disequilibrium in GWAS and downstream applications (Nowbandegani et al., 2023).

ARG-based methods can simulate genetic variation for millions of samples and store the output very compactly in the “succinct tree sequence” encoding (Wong et al., 2023) and `tskit` library (Ralph et al., 2020). For example, a highly realistic simulation of chromosome 9 for 1.4 million French-Canadian samples (Anderson-Trocmé et al., 2023) requires around 550GB of storage space in gzip-compressed VCF (Danecek et al., 2011). At 1.36GB the original simulated ARG (compressed using the `tszip` utility) is around 400X smaller. Furthermore, many calculations can be expressed efficiently in terms of the underlying ARG (Kelleher et al., 2016; Ralph et al., 2020), without needing to decode the actual variation data. Finally, outputting a simulated ARG provides access to the full history, not just the genetic variation among the samples.

Although there are sophisticated methods available for simulating ARGs at biobank scale, there is currently no easy way to simulate quantitative traits based on such an ARG. Many existing methods to simulate quantitative traits from a given set of genetic sequences assume that the genotypes fit in memory (e.g. Meyer and Birney, 2018; Fernandes and Lipka, 2020), which makes them impractical at biobank scale (the French-Canadian dataset discussed above would require 140TB of RAM assuming 1 byte per genotype). Methods that read parts of the genotype matrix from file as required (e.g. Wharrie et al., 2023) can be used on reasonable hardware, but working with such large files is slow and cumbersome. More fundamentally, exporting genotypes discards much of rich detail about ancestral history contained in an ARG, and it is exactly this information that we wish to take advantage of when using inferred ARGs in GWAS applications. In their analysis of the portability of polygenic risk scores across populations, Martin et al. (2017) demonstrated the utility of simulating phenotypes directly from an ARG. Their approach, however, is tightly coupled to the details of the study and not designed to be reused. Simulation code can be subtle and difficult to debug (Ragsdale et al., 2020), and there is a critical need for a well-documented and thoroughly tested means of simulating quantitative traits directly from an ARG.

In this paper we present `tstrait`, a Python library that efficiently simulates quantitative traits on an arbitrary ARG. `Tstrait` can quickly simulate quantitative traits for population-scale datasets, with a very low memory overhead, and taking into account the rich historical detail contained within an ARG. The `tstrait` library also integrates well with the wider Python data-science ecosystem (Harris et al., 2020), allowing users to efficiently analyse large-scale data using familiar and ergonomic tools.

## Results

### Model

Phenotypes are simulated in `tstrait` following standard GWAS models (Uffelmann et al., 2021), adapted to the ARG context. Each trait is associated with one or more causal sites (positions on the genome), and at each causal site there is a causal allele (i.e. a particular nucleotide) associated with an effect size *β*. For each causal site, an effect size is drawn from a distribution and optionally multiplied by (2*p*(1 *− p*))^*α/*2^, where *p* is the frequency of the causal allele and *α* is a parameter describing the strength of frequency dependence (Speed et al., 2012). At a particular causal site, every node in the local tree that inherits the causal allele at that site is said to have a “genetic value” of *β*.

In Fig. 1 we show an example tree for three individuals. Because these individuals are diploids, each is associated with two nodes in the tree (highlighted by colour). Ancestral nodes are not associated with individuals here, but in general an ARG may be embedded in a multigenerational pedigree, where some internal nodes would be associated with individuals. In the example of Fig. 1, T is chosen as the causal allele with *β* = 0.05, so all nodes descending from i have genetic value 0.05, except e which has zero because of the back-mutation to A. Following the standard practise in GWAS (Uffelmann et al., 2021), we assume the additive model such that the overall genetic value of an individual is the sum of its nodes’ genetic values. Given these per-individual genetic values, the final phenotype is then generated by adding some environmental noise. This noise is simulated from a normal distribution with mean zero and variance of *V*_*G*_(1 *− h*^2^)*/h*^2^, where *V*_*G*_ is the variance of the individual genetic values and *h*^2^ is the narrow-sense heritability provided as input by the user.

**Fig. 1:**
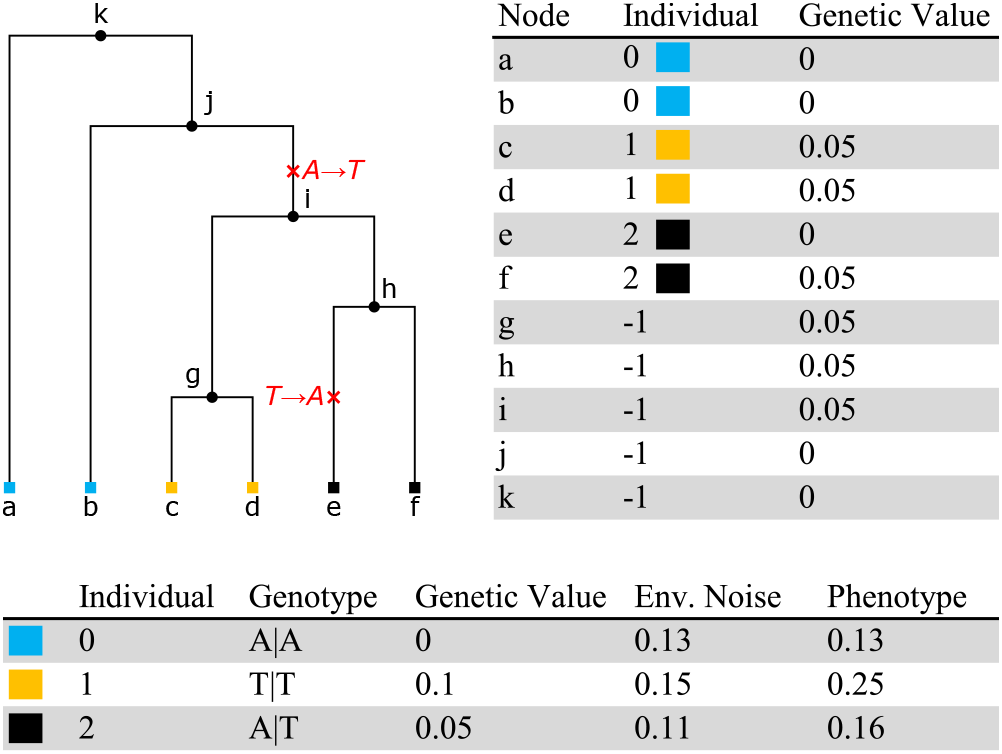
Example simulation of a phenotype at a site with ancestral state A and two mutations. In this diploid example each of the three individuals is associated with two nodes (i.e., the individual with ID 0 corresponds to nodes a and b). Internal nodes in the tree are associated with the null individual, *−*1. Here, the trait’s causal allele is T with an effect size *β* = 0.05. Each node in the tree has an associated genetic value, and the overall genetic value for an individual is the sum of the genetic values of their corresponding nodes. The final phenotype for each individual is the sum of the genetic value and simulated environmental noise.

### Interface

`Tstrait` is a Python library, building on the `tskit` ARG toolkit (Ralph et al., 2020; Wong et al., 2023) and the rich Python data-science ecosystem (Harris et al., 2020). Simulating a phenotype for an ARG with default parameter values requires only a few lines of code:

~~~
**import** tstrait as tst
model = tst. trait_model (‘normal’, mean=0, var =1)
result = tst. sim_phenotype (arg, model)
~~~

We first create `model`, representing the distribution from which effect sizes are drawn. Five commonly used univariate distributions are supported, along with the multivariate normal distribution to model pleiotropic traits. Given this model, we can then simulate phenotypes for the individuals in an ARG (as a `tskit TreeSequence`) using the `sim_phenotype` function. The user can either specify a number of causal sites to be chosen randomly along the genome (one, by default), or can directly provide the causal sites as input. Combined with the detailed information about mutations recorded in a `tskit` ARG, explicitly specifying causal sites allows us to model many different types of trait, for example those associated with mutations arising in a particular population or time interval. The return value `result` is an object encapsulating two Pandas dataframes (McKinney, 2010): one describing the simulated effect sizes and the other describing the genetic values, environmental noise and phenotypes for each individual. The simulation results can then be efficiently and conveniently processed using standard Python data science tools.

As well as this convenient single-function interface, `tstrait` provides modular building blocks for power-users and to facilitate integration with other tools that generate traits on an ARG. The `sim trait` function simulates effect sizes for an input ARG, and returns a data frame describing the causal sites, alleles and effect sizes. This dataframe can then be passed to the `genetic_values` function, which calculates the genetic values for each node, and accumulates them by individual (Fig 1). Finally, the `sim env` function takes these per-individual genetic values and adds some simulated environmental noise to produce the final phenotypes.

A major benefit of this modular architecture is the flexibility it offers users. Because the causal sites and effect sizes are specified in a simple tabular format, users can easily develop their own approach to simulating these values. Alternatively, other simulators such as SLiM (Haller and Messer, 2023) that generate effect sizes and causal mutations during the progress of a forwards-time simulation could output these values to a CSV or similar file. The modular architecture and simple input data formats are specifically intended to facilitate such interoperability.

### Implementation and validation

`Tstrait` is written entirely in Python. Numerical operations are either peformed using standard array-oriented approaches (Harris et al., 2020) or accelerated using the `numba` JIT compiler (Lam et al., 2015). The `tstrait` codebase includes a suite of unit tests, which are automatically run as part of the development process. The output of `tstrait` has been validated against theoretical expectations, as well as the output of `AlphaSimR` (Gaynor et al., 2021) and `simplePHENOTYPES` (Fernandes and Lipka, 2020).

### Performance

`Tstrait` is very efficient, and can be applied to datasets at the largest scales on standard computers. Supplementary Fig S1 shows how trait simulation time scales with number of individuals on human-like coalescent simulations generated using `stdpopsim` (Adrion et al., 2020). To emphasise scalability, we also applied `tstrait` to the large French-Canadian simulations discussed in the introduction. It took 80.69 seconds to simulate a trait with 100 causal sites for all 2.7 million pedigree individuals. Finally, to demonstrate that `tstrait` can also be applied to ARGs inferred from real data, we simulated a trait with 100 causal sites for an ARG estimated from 1000 Genomes project data (Kelleher et al., 2019) which has 2,504 samples and 1,685,401 variant sites. This took 5.40 seconds. Memory requirements for `tstrait` are modest: all of the above experiments were performed on a laptop computer with 16GB of RAM.

## Conclusion

There is substantial interest in using inferred ARGs to improve association testing methods (Zhang et al., 2023; Link et al., 2023; Nowbandegani et al., 2023), and there is a pressing need for a well-tested, efficient and user-friendly means of simulating phenotypes on ARGs. Highly realistic simulations conditioned on large pedigrees (Anderson-Trocmé et al., 2023) provide an exciting opportunity to test the effects of intricate population structure on GWAS, and we hope that `tstrait` will facilitate these investigations. `Tstrait`’s modular architecture and flexible specification of causal sites should provide the opportunity to explore new avenues of research, and an extensible platform for future development.

## Acknowledgments

We are grateful to Gregor Gorjanc, Ben Haller, Ben Jeffery, Pier Palamara, Alison Etheridge and others in the tskit community for helpful discussions and feedback.

## Supplementary Figures

**Fig. S1:**
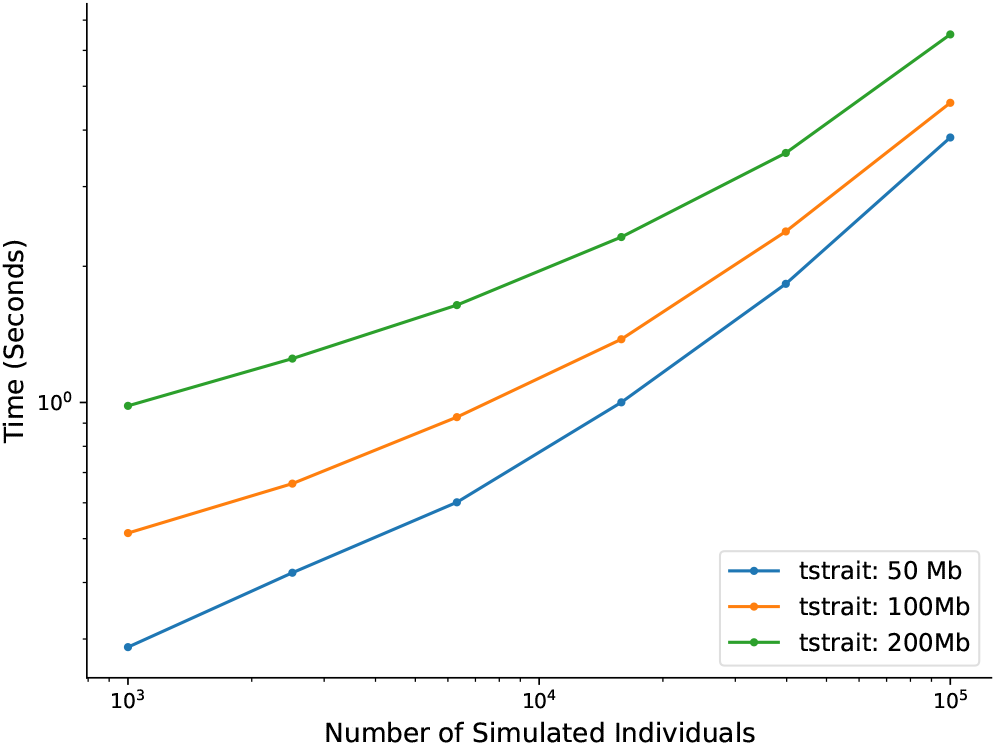
Time taken to simulate quantitative traits with increasing sample size. For each sample size, we simulated an ARG under human-like parameters using the default HomSap demographic model in `stdpopsim`. Each point represents the mean time for 10 independent runs of `tstrait` for a particular ARG. The times reported are the total CPU time required to simulate a quantitative trait with 1000 causal sites, on an Intel(R) Core(TM) i9-11900H CPU and 16 GB of RAM. The trait model is a normal distribution with *μ* = 0, *σ*^2^ = 1, *h*^2^ = 0.3, and *α* = 0.

